# Genomic Correlates of APOBEC-Associated Mutagenesis in Esophageal Squamous Cell Carcinoma

**DOI:** 10.64898/2026.10.27.762354

**Authors:** Emmanuel Owusu Ansah, Joshua Ampofo Yentumi, Emmanuel Yeboah Kodom, Prince Owusu Mensah, Kwadwo Fosu, Isaac Kofi Adu, Emmanuel Adu Boahen

## Abstract

**Background and Aims:** APOBEC-associated mutagenesis is prominent in esophageal squamous cell carcinoma (ESCC), but genomic features associated with its intertumoral variability remain incompletely characterized. We investigated genomic correlates of APOBEC-associated mutagenesis while accounting for the compositional structure of mutational signature data.

**Methods:** Mutational signatures were analyzed in 552 ESCC tumors from eight countries. APOBEC-associated mutagenesis was quantified as SBS2 plus SBS13 using absolute mutation counts and proportional contributions. Associations with co-occurring mutational signatures, total SBS burden, recurrent driver-gene alterations, and country were evaluated using correlation, sensitivity, and multivariable analyses. Principal component and regression-model robustness were additionally assessed.

**Results:** APOBEC burden varied markedly across tumors (median, 2,002 mutations; IQR, 987–4,346). Several associations observed using signature proportions were not maintained in absolute-count analyses: SBS1, SBS3, and SBS5 changed direction, indicating sensitivity to compositional representation. In contrast, SBS18 remained inversely associated with APOBEC-associated mutagenesis in all variable pairings examined: proportion versus proportion (Spearman ρ = −0.527), APOBEC proportion versus SBS18 absolute count (r = −0.435), and absolute count versus absolute count (Pearson r = −0.319, P = 1.59 × 10□^1^□; Spearman ρ = −0.317). These are distinct comparisons rather than discrepant estimates. Notably, this cohort-wide relationship was primarily characterized by higher APOBEC burden in SBS18-negative than SBS18-positive tumors rather than a continuous inverse relationship among SBS18-positive tumors. The multivariable model had an in-sample R^2^ of 0.400 but lower and variable 10-fold cross-validated performance (mean R^2^ = 0.273). Country added little explanatory information to this model (ΔR^2^ = 0.008; global P = 0.402). ZNF750-mutant tumors had higher APOBEC burden than wild-type tumors (BH-FDR q = 0.0032), although only 12 tumors carried ZNF750 alterations.

**Conclusion:** Genomic correlates of APOBEC-associated mutagenesis in ESCC are sensitive to the representation of mutational signature activity. SBS18 showed the most consistent inverse association, whereas the ZNF750 association warrants independent validation.

## Introduction

Esophageal squamous cell carcinoma (ESCC) is one of the leading causes of cancer-related mortality worldwide [1]. It remains a major global health burden, particularly in regions of Asia and Africa [2]. Despite substantial geographic variation in incidence [3], genomic studies have shown that the overall mutational landscape of ESCC is broadly conserved across populations [4,5]. Large-scale sequencing studies have identified several recurrent mutational processes in ESCC, including age-related and clock-like signatures as well as endogenous DNA-editing processes [5]. Among these, mutagenesis associated with the apolipoprotein B mRNA editing enzyme catalytic polypeptide-like (APOBEC) family is a prominent feature of ESCC genomes [6,7].

The COSMIC single-base substitution signatures SBS2 and SBS13 are widely recognized as mutational footprints of APOBEC-mediated cytidine deamination and have been detected across multiple cancer types [8]. These signatures arise from the deamination of cytosine in single-stranded DNA, followed by distinct downstream processing that produces characteristic mutation patterns. In ESCC, SBS2 and SBS13 are detected in a substantial proportion of tumors and can contribute considerably to overall mutation burden [5]. Previous studies have linked APOBEC-associated mutagenesis to biological processes including replication stress, immune signaling, DNA repair, and genomic instability, suggesting that APOBEC-mediated mutagenesis may contribute to tumor evolution and genomic diversification [9–11].

Despite its prominence in ESCC, the genomic features associated with intertumoral variation in APOBEC-associated mutagenesis remain incompletely characterized. Established or proposed biological determinants of APOBEC mutagenesis include germline variation involving *APOBEC3A* and *APOBEC3B* [12], differences in APOBEC gene expression [13], and differences in the mutagenic contributions of individual APOBEC enzymes [14]. However, these factors do not fully describe the broader genomic context in which APOBEC-associated mutations accumulate. Examining relationships between APOBEC-associated mutagenesis and other tumor genomic features may therefore provide insight into the heterogeneity of this mutational process across ESCC tumors.

An additional challenge is that mutational signature activity can be represented either as absolute mutation counts or as proportions of the total mutational burden. Signature proportions are compositional because the contributions of all signatures within a tumor are constrained by their total. Consequently, an increase in the proportional contribution of one signature necessarily changes the relative contributions of other signatures, potentially producing associations that do not correspond to relationships between their absolute mutation counts [15–17]. This issue is particularly relevant when a mutational process such as APOBEC-associated mutagenesis accounts for a substantial fraction of mutations in some tumors. Evaluating associations using both proportional and absolute representations is therefore important for distinguishing robust genomic relationships from those that may depend on the statistical representation of mutational signature activity.

In this study, we performed a secondary analysis of whole-genome sequencing data from 552 ESCC tumors from eight countries to characterize genomic correlates of APOBEC-associated mutagenesis, quantified using SBS2 and SBS13. We examined its relationships with total single-base substitution burden, co-occurring mutational signatures, recurrent driver-gene alterations, and geographic origin using continuous proportional and absolute-count analyses. We further assessed the robustness of these relationships through sensitivity analyses addressing the compositional structure of mutational signature data, principal component analysis excluding SBS2 and SBS13, regression diagnostics and internal validation, and multiple-testing-adjusted analysis of prespecified driver genes. The objective was to identify genomic features associated with intertumoral variation in APOBEC-associated mutagenesis while distinguishing statistical associations from evidence of biological or causal determinants.

## Materials and Methods

### Ethical Approval

This study consisted of a secondary analysis of de-identified, previously generated whole-genome sequencing data obtained from the European Genome–Phenome Archive (EGA, accession EGAD00001006868). Filtered somatic variant call files used for signature analysis were obtained from the companion dataset EGAD00001008339, released under the same parent study (EGAS00001002725). Ethical approval and participant consent for sample and data collection were obtained by the original investigators as part of the primary international ESCC sequencing study [5]. As the present analysis used only de-identified, aggregated mutational signature data with no additional patient contact, no separate institutional ethical approval was required for this secondary analysis, in accordance with institutional policy.

### Overview of the Study Design

The primary objective of this study was to characterize genomic features associated with intertumoral variability in APOBEC-associated mutagenesis in esophageal squamous cell carcinoma (ESCC). We evaluated relationships between APOBEC-associated mutagenesis, overall single-base substitution (SBS) burden, co-occurring mutational signatures, recurrent driver-gene alterations, and country of origin in an international cohort of ESCC tumors. Analyses were performed using tumor-level mutational signature profiles derived from whole-genome sequencing data. APOBEC-associated mutagenesis was evaluated as a continuous quantitative trait using both absolute mutation counts and relative contributions to the total SBS burden. Because mutational signature proportions are compositional and therefore subject to constant-sum dependence, analyses based on absolute signature counts were used to assess the robustness of associations observed using proportional measures. Correlation analyses, principal component analysis (PCA), multivariable regression, driver-gene analyses, and geographic sensitivity analyses were performed. Median-based APOBEC groups were retained only for descriptive visualization and were not used as the primary basis for inference.

### Study Cohort and Data Sources

The study included 552 ESCC tumors from a previously published international whole-genome sequencing cohort [5]. Tumors were obtained from eight countries representing regions with differing ESCC incidence: Brazil, China, Iran, Japan, Kenya, Malawi, Tanzania, and the United Kingdom. Country of origin and associated study information were obtained from the accompanying metadata and study documentation. Somatic SBS mutations were represented as tumor-level activities assigned to COSMIC mutational signatures. The signatures identified in the analyzed dataset were SBS1, SBS2, SBS3, SBS5, SBS8, SBS9, SBS13, SBS15, SBS17a, SBS17b, SBS18, SBS33, SBS44, and SBS102. All analyses were conducted at the tumor level.

### Mutational Signature Analysis

SBS96 mutation matrices were generated from somatic variant data using SigProfilerMatrixGenerator v1.3.6 with the GRCh37 reference genome. De novo mutational signature extraction was performed using SigProfilerExtractor v1.2.6, with one to five signatures evaluated using non-negative matrix factorization. Extracted signatures were decomposed against the COSMIC v3.5 SBS reference catalogue for GRCh37, and signature activities were assigned using SigProfilerAssignment v1.1.3. The resulting absolute signature activities, expressed as the numbers of mutations assigned to each signature in each tumor, were used for downstream analyses. SBS102 was retained in the analyzed signature profiles because it was present in the output generated by the signature-assignment workflow and was not introduced manually. Relative signature contributions were calculated by dividing individual signature activities by the total SBS mutation count for each tumor.

### Quantification of APOBEC-Associated Mutagenesis

APOBEC-associated mutagenesis was defined as the combined contribution of SBS2 and SBS13, which are associated with APOBEC cytidine deaminase activity [8]. For each tumor, absolute APOBEC burden was calculated as the number of mutations assigned to SBS2 plus SBS13. APOBEC proportion was calculated as the absolute APOBEC burden divided by the total number of SBS mutations.

Total SBS burden was calculated as the sum of the activities of all identified SBS signatures. For analyses requiring a measure of mutation burden independent of APOBEC-associated mutations, non-APOBEC background burden was calculated as total SBS burden minus the SBS2 and SBS13 mutation counts. Both absolute and proportional measures were examined because signature proportions constitute compositional data. Associations observed using proportions were therefore compared with analyses based on absolute mutation counts to distinguish relationships that were robust to representation from those potentially arising from constant-sum dependence.

### Association Between APOBEC-Associated Mutagenesis and Mutational Signatures

Associations between APOBEC-associated mutagenesis and individual non-APOBEC mutational signatures were evaluated using Pearson and Spearman correlation coefficients. For SBS18, four pairings were reported separately and labelled throughout: proportion versus proportion, APOBEC proportion versus SBS18 absolute count, absolute count versus absolute count, and absolute count versus absolute count with adjustment for non-APOBEC background burden (partial correlation). Primary sensitivity analyses compared correlations based on absolute APOBEC and signature mutation counts with corresponding correlations based on proportional signature contributions. P values were adjusted across tested signatures using the Benjamini–Hochberg false discovery rate (FDR) procedure.

Because SBS18 showed the most consistent inverse relationship with APOBEC-associated mutagenesis, additional analyses were performed to characterize this association. Tumors were classified according to the presence or absence of detectable SBS18 activity, and absolute APOBEC burdens were summarized within these groups. The association between absolute SBS18 and APOBEC mutation counts was also evaluated separately among SBS18-positive tumors. Additional regression analyses examined the APOBEC–SBS18 relationship while accounting for background mutation burden. These analyses were used to distinguish an overall cohort-level association from a continuous relationship among tumors with detectable SBS18 activity.

### Principal Component Analysis

PCA was performed using tumor-level absolute mutational signature activities without feature standardization. In the initial analysis, all identified SBS signatures were included, and the proportion of variance explained and signature loadings for the principal components were recorded. Pearson and Spearman correlations were used to describe associations between APOBEC proportion and principal-component scores.

Because SBS2 and SBS13 directly define the APOBEC measure, associations between APOBEC-associated mutagenesis and principal components derived from a matrix containing these signatures are not statistically independent. The original PCA was therefore treated as descriptive. As a sensitivity analysis, PCA was repeated after excluding SBS2 and SBS13 from the signature matrix. Associations between APOBEC-associated mutagenesis and the resulting principal components were then reevaluated to determine whether the relationships persisted when the signatures used to define APOBEC-associated mutagenesis were not themselves used to construct the principal components.

### Multivariable Regression and Model Diagnostics

A multivariable ordinary least-squares regression model was fitted with APOBEC proportion as the dependent variable and total SBS burden, SBS1, SBS3, SBS5, and SBS18 absolute mutation counts as explanatory variables. Heteroskedasticity-consistent HC3 standard errors were used for statistical inference. Model fit was summarized using the coefficient of determination (R^2^) and adjusted R^2^. Potential numerical multicollinearity was assessed using variance inflation factors (VIFs) and condition indices calculated from standardized predictor variables. Coefficient stability was further examined by sequentially removing individual predictors and comparing the resulting coefficients with those from the full model.

Internal model performance was assessed using 10-fold cross-validation with reproducible randomized fold assignment. R^2^ was calculated for each held-out fold and summarized by the mean, median, and standard deviation across folds. Bootstrap resampling with 2,000 replicates was additionally used to characterize the distributions of model R^2^, adjusted R^2^, and regression coefficients. Because total SBS burden contains the APOBEC-associated mutations used to construct the outcome proportion, this model was interpreted as an associative description of the observed signature structure rather than as a causal or predictive model.

### Geographic Analyses

Differences in APOBEC proportion across the eight countries were first evaluated descriptively and using the Kruskal–Wallis test. To examine whether country contributed additional information after accounting for the genomic variables included in the multivariable model, country was subsequently added as a categorical covariate, with Brazil specified as the reference category. The contribution of country as a whole was evaluated using a nested-model F test comparing models with and without the seven country indicator variables. Change in model R^2^ (ΔR^2^) was also calculated. In addition, country-stratified sensitivity analyses evaluated the relationship between absolute APOBEC and SBS18 mutation counts within each country.

Because potentially relevant individual-level covariates, including age, smoking, alcohol use, and tumor purity, were not available in the metadata used for this secondary analysis, and biological sex was unavailable for most tumors, the country-adjusted analyses were considered exploratory rather than evidence excluding geographic or exposure-related influences.

### APOBEC-Based Tumor Stratification

For descriptive analyses and visualization, tumors were classified as APOBEC-high or APOBEC-low using the cohort median APOBEC proportion as the threshold. This stratification was used to display differences in mutational composition and to facilitate comparison with the original analyses. Because the groups were defined directly from APOBEC proportion, group-based comparisons of complementary signature proportions were not treated as independent evidence of associations between mutational processes. Inferential conclusions were instead based primarily on continuous analyses and corresponding absolute-count sensitivity analyses.

### Driver-Gene Analysis

Protein-altering alterations in 11 established ESCC driver genes—*TP53, KMT2D, CDKN2A, NOTCH1, PIK3CA, FAT1, NFE2L2, KEAP1, ZNF750, CUL3*, and *AJUBA*—were identified from annotated variant call files and represented as binary mutation-status variables. For each gene, absolute APOBEC mutation counts were compared between mutant and wild-type tumors using the Mann–Whitney U test, with rank-biserial correlation used as an effect-size measure. Benjamini–Hochberg FDR correction was applied across the 11 genes. As a complementary analysis, log-transformed APOBEC burden was modeled as a function of driver-gene mutation status while adjusting for non-APOBEC background mutation burden, defined as total SBS burden minus SBS2 and SBS13 counts. Heteroskedasticity-consistent HC3 standard errors were used, and FDR correction was applied across the 11 driver-gene tests. Concordance between the rank-based and adjusted analyses was considered when interpreting driver-gene associations. For *ZNF750*, individual protein-altering variants were further characterized by mutation class and local sequence context. Sequence contexts were compared with canonical SBS2- and SBS13-compatible motifs to assess whether *ZNF750* mutations themselves were consistent with direct APOBEC-associated mutagenesis. Given the small numbers of tumors carrying several driver alterations, these analyses were considered hypothesis-generating.

### Statistical Analysis and Reproducibility

Analyses were conducted in Python 3.13.2. Major software packages included pandas v2.3.2, NumPy v2.3.2, SciPy v1.16.2, scikit-learn v1.8.0, statsmodels, and Matplotlib. Mutational signature analyses used SigProfilerMatrixGenerator v1.3.6, SigProfilerExtractor v1.2.6, SigProfilerAssignment v1.1.3, and SigProfilerPlotting v1.4.3. All statistical tests were two-sided. Unless otherwise specified, P < 0.05 was considered statistically significant, and the Benjamini–Hochberg procedure was used to control the FDR for families of multiple comparisons. Analyses were implemented using reproducible Python scripts, and a single locked tumor-level dataset was used for the revised analyses to ensure consistency across tables, figures, and statistical models. The analysis code is publicly available (see Code availability).

## Results

### Distribution of APOBEC-associated mutagenesis in ESCC

Across the 552 ESCC tumors, absolute APOBEC-associated mutation burden, defined as th combined number of SBS2 and SBS13 mutations, ranged from 0 to 43,883 mutations, with a median of 2,002 (IQR, 987–4,346) and a mean of 3,764.5. APOBEC-associated mutations accounted for a median of 20.3% of total SBS mutations (IQR, 12.1%–33.8%). Forty-six tumors (8.3%) had no detectable SBS2 or SBS13 mutations (Table 1). Tumor-level mutational signature activities and genomic characteristics are provided in Supplementary Table S1. APOBEC proportion was positively associated with total SBS burden (Pearson r = 0.396, P = 3.79 × 10□^22^; Spearman ρ = 0.368, P = 3.87 × 10□^1^□; Figure 3A), indicating that tumors with a greater overall mutational burden tended to have a higher relative contribution from SBS2 and SBS13.

**Table 1.** Cohort characteristics and APOBEC-associated mutagenesis summary, 552 ESCC tumors.

| **Characteristic** | **Value** |
|---|---|
| Tumors, N | 552 |
| Total SBS mutations, median (IQR) | 10,709 (7,888–15,347) |
| APOBEC-associated mutations (SBS2 + SBS13), median (IQR) | 2,002 (987–4,346) |
| APOBEC-associated mutations, mean (SD) | 3,764.5 (5,152.9) |
| APOBEC proportion, median (IQR) | 0.203 (0.121–0.338) |
| APOBEC proportion, mean (SD) | 0.253 (0.186) |
| Tumors with no SBS2/SBS13 mutations, n (%) | 46 (8.3%) |
| SBS18 absent, n (%) | 208 (37.7%) |
| SBS18 present, n (%) | 344 (62.3%) |
| Country: Iran, n (%) | 178 (32.2%) |
| Country: China, n (%) | 138 (25.0%) |
| Country: Kenya, n (%) | 68 (12.3%) |
| Country: Malawi, n (%) | 59 (10.7%) |
| Country: Japan, n (%) | 37 (6.7%) |
| Country: Tanzania, n (%) | 35 (6.3%) |
| Country: Brazil, n (%) | 30 (5.4%) |
| Country: UK, n (%) | 7 (1.3%) |

### Multivariate structure of mutational signatures

In the original PCA including all mutational signatures, PC1 explained 36.2% of the total variance and was strongly associated with APOBEC proportion (Pearson r = 0.759, P = 1.83 × 10□^1^□□; Figure 3C). SBS13 and SBS2 contributed strongly to PC1, with loadings of 0.815 and 0.487, respectively. Because these signatures also define the APOBEC measure, we performed a sensitivity PCA after excluding SBS2 and SBS13. In this analysis, PC1 explained 31.3% of the remaining variation and was no longer associated with APOBEC proportion (Pearson r = 0.010, P = 0.806; Spearman ρ = −0.015, P = 0.733; Supplementary Figure S1; Supplementary Table S2). Thus, the strong association between APOBEC proportion and PC1 in the original PCA was largely attributable to the direct contribution of SBS2 and SBS13 to the principal component.

### Associations between APOBEC-associated mutagenesis and other mutational signatures

Several mutational signatures showed associations with APOBEC proportion, although the direction and magnitude of these relationships differed when absolute mutation counts were considered (Figure 2; Table 2). The strongest inverse association was observed for SBS18. Four related but distinct SBS18 estimates are reported and should not be conflated. (i) Proportion versus proportion: SBS18 proportion was inversely correlated with APOBEC proportion (Pearson r = −0.535; Spearman ρ = −0.527; Table 2). (ii) APOBEC proportion versus SBS18 absolute count: r = −0.435, P = 7.90 × 10□^2^□ (Figure 3B). (iii) Absolute count versus absolute count: Pearson r = −0.319, P = 1.59 × 10□^1^□; Spearman ρ = −0.317, P = 2.19 × 10□^1^□ (Figure 4A; Table 2). (iv) Absolute count versus absolute count after adjustment for non-APOBEC background mutation burden: partial Pearson r = −0.290, P = 3.44 × 10□^12^. Because the absolute-count estimates are least affected by the constant-sum constraint, they are used as the principal measure of the relationship, and the inverse association therefore persisted when absolute mutation counts were analyzed. In contrast, SBS1, SBS3, and SBS5 showed positive correlations with absolute APOBEC burden despite inverse relationships when expressed as proportions (Table 2), highlighting the influence of compositional scaling on proportional signature analyses.

**Table 2.** Associations between APOBEC-associated mutagenesis and individual mutational signatures, using both absolute mutation counts and proportional contributions. “Direction” indicates whether the sign of the association is the Same or Opposite between the two representations. For SBS18, coefficients in the proportional columns refer to SBS18 proportion versus APOBEC proportion, and coefficients in the absolute columns refer to SBS18 count versus APOBEC count; the APOBEC proportion versus SBS18 count estimate (r = −0.435) shown in Figure 3B is a separate comparison.

| Signature | Count r (Pearson) | Count FDR | Count ρ (Spearman) | Count FDR | Prop. r (Pearson) | Prop. FDR | Prop. ρ (Spearman) | Prop. FDR | Direction |
|---|---|---|---|---|---|---|---|---|---|
| SBS1 | 0.202 | 6.83e-06 | 0.398 | 2.69e-21 | -0.312 | 2.72e-13 | -0.245 | 2.23e-08 | Opposite |
| SBS3 | 0.178 | 7.85e-05 | 0.104 | 0.035 | -0.133 | 0.004 | -0.102 | 0.020 | Opposite |
| SBS5 | 0.273 | 4.12e-10 | 0.326 | 2.20e-14 | -0.347 | 3.06e-16 | -0.316 | 1.57e-13 | Opposite |
| SBS8 | -0.080 | 0.148 | -0.135 | 0.004 | -0.171 | 1.61e-04 | -0.166 | 2.73e-04 | Same |
| SBS9 | -0.063 | 0.277 | -0.070 | 0.163 | -0.129 | 0.005 | -0.108 | 0.018 | Same |
| SBS15 | -0.029 | 0.657 | -0.045 | 0.390 | -0.076 | 0.082 | -0.104 | 0.020 | Same |
| SBS17a | -0.029 | 0.657 | -0.026 | 0.595 | -0.087 | 0.054 | -0.065 | 0.140 | Same |
| SBS17b | 0.015 | 0.796 | 0.011 | 0.800 | -0.084 | 0.059 | -0.041 | 0.334 | Opposite |
| SBS18 | -0.319 | 1.91e-13 | -0.317 | 8.77e-14 | -0.535 | 4.32e-41 | -0.527 | 1.01e-39 | Same |
| SBS33 | 0.007 | 0.861 | -0.034 | 0.516 | -0.046 | 0.278 | -0.107 | 0.018 | Opposite |
| SBS44 | -0.020 | 0.766 | -0.068 | 0.163 | -0.108 | 0.020 | -0.117 | 0.012 | Same |
| SBS102 | -0.045 | 0.501 | -0.095 | 0.051 | -0.104 | 0.022 | -0.131 | 0.005 | Same |

**Fig. 1.**
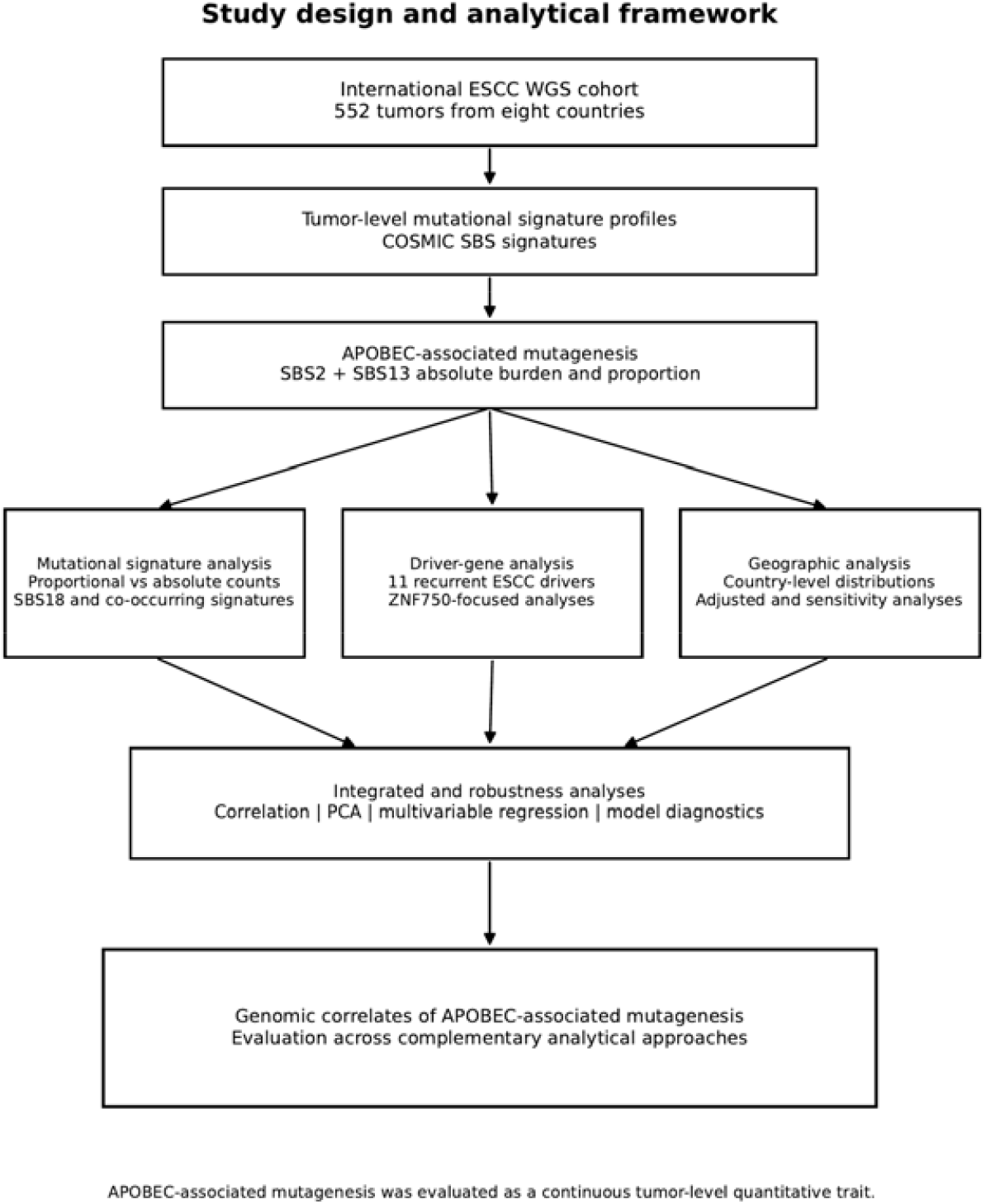
Study design and analytical framework. Whole-genome sequencing data from 552 ESCC tumors across eight countries were used to derive tumor-level mutational signature profiles against the COSMIC SBS reference. APOBEC-associated mutagenesis was quantified as the SBS2 + SBS13 absolute burden and proportion and evaluated through three complementary analytical branches: mutational signature analysis (proportional versus absolute counts, SBS18 and co-occurring signatures), driver-gene analysis (11 recurrent ESCC driver genes, with ZNF750-focused analyses), and geographic analysis (country-level distributions with adjusted and sensitivity analyses). These branches were integrated through correlation, principal component analysis, multivariable regression, and model diagnostics to characterize genomic correlates of APOBEC-associated mutagenesis across complementary analytical approaches, with APOBEC-associated mutagenesis modeled throughout as a continuous tumor-level quantitative trait.

**Fig. 2.**
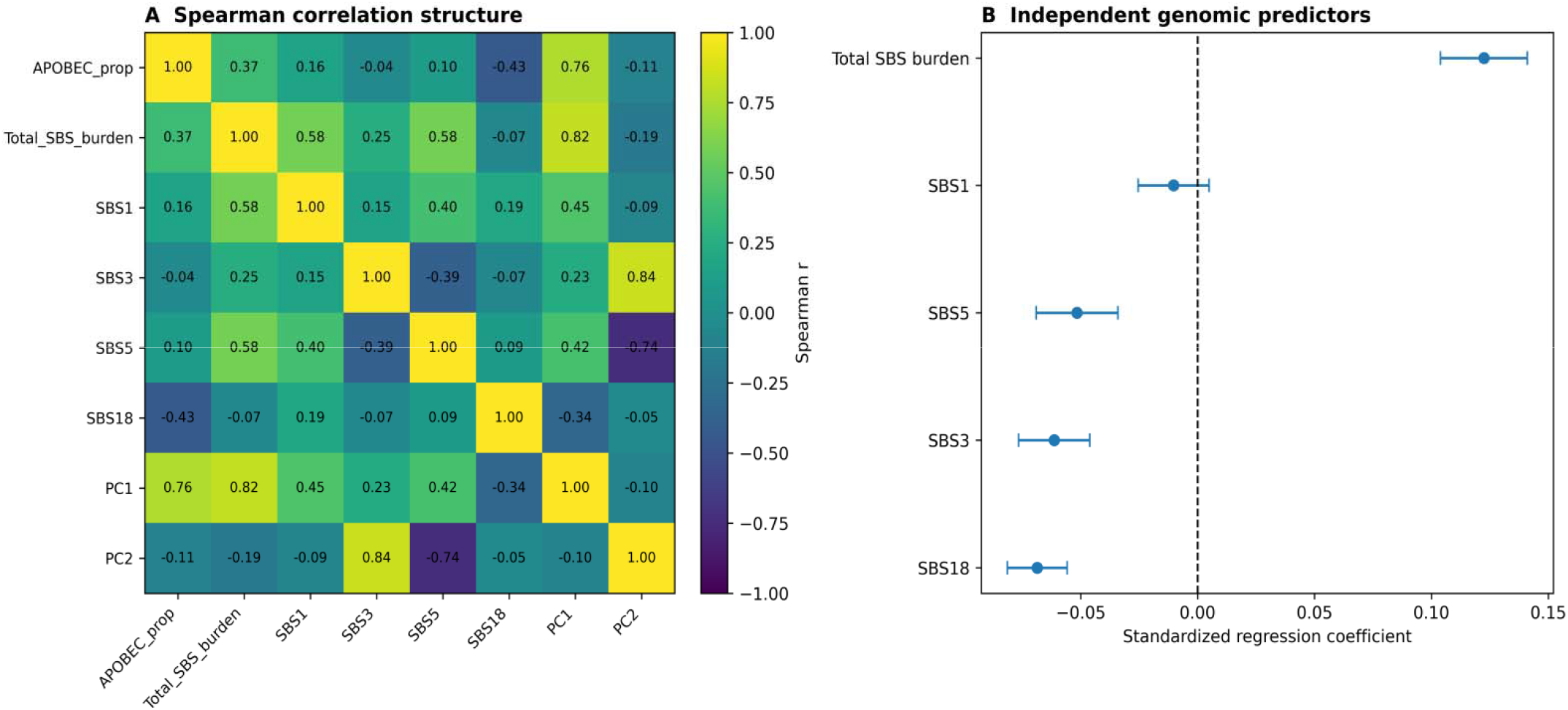
Correlation structure and independent genomic predictors of APOBEC-associated mutagenesis in ESCC. (A)Spearman correlation heatmap showing relationships between APOBEC-associated mutagenesis, total SBS burden, major mutational signatures, and principal components. (B)Standardized regression coefficients from multivariable linear regression analysis identifying independent genomic predictors of APOBEC-associated mutagenesis. Error bars represent 95% confidence intervals.

**Fig. 3.**
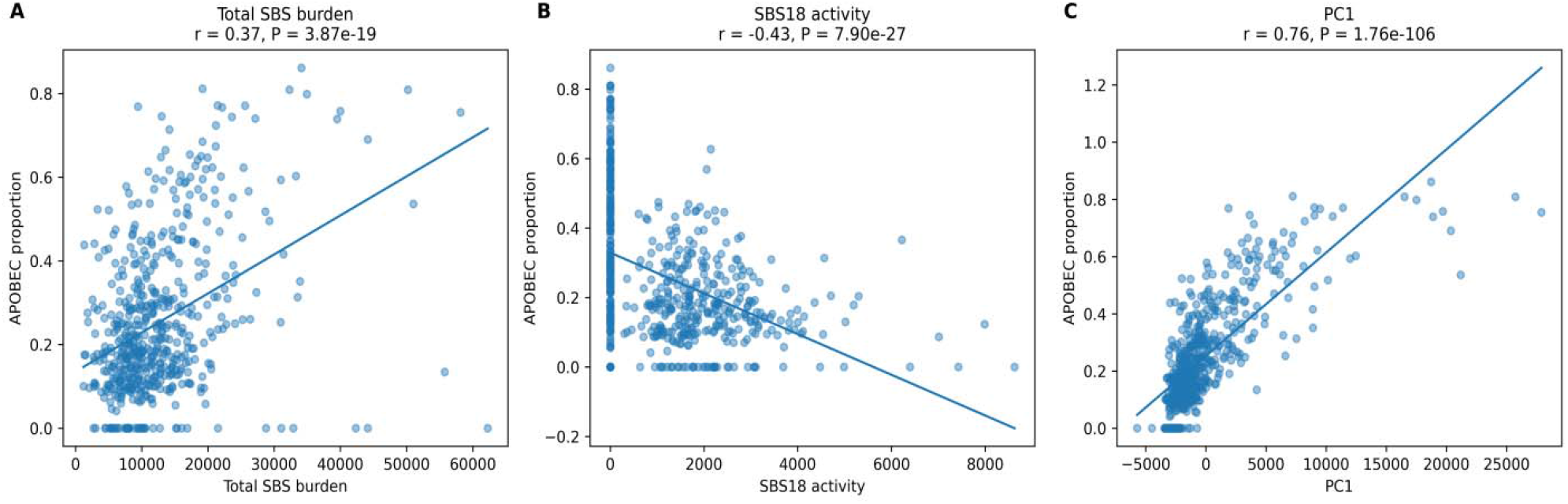
Key genomic relationships associated with APOBEC activity in ESCC tumors. (A)Association between APOBEC-associated mutagenesis and total SBS mutation burden. (B)Inverse association between APOBEC-associated mutagenesis and SBS18 (APOBEC proportion versus SBS18 absolute count; r = −0.435). (C)Association between APOBEC-associated mutagenesis and PC1 derived from mutational signature profiles. Spearman correlation coefficients and corresponding P values are shown.

Further examination of SBS18 showed that 208 tumors had no detectable SBS18, whereas 344 had detectable SBS18. Median absolute APOBEC burden was 4,638.5 mutations in SBS18-absent tumors compared with 1,605.5 mutations in SBS18-positive tumors (Figure 4B). Among SBS18-positive tumors, however, absolute SBS18 and APOBEC mutation counts were positively rather than negatively correlated (Pearson r = 0.163, P = 0.0024; Spearman ρ = 0.266, P = 5.75 × 10□□). After adjustment for genomic covariates, the SBS18-presence effect was not significant (β = −0.107, 95% CI −0.821 to 0.607, P = 0.769). Together, these findings indicate that the cohort-wide inverse relationship was strongly influenced by differences between SBS18-absent and SBS18-positive tumors rather than a simple inverse relationship among tumors with detectable SBS18.

**Fig. 4.**
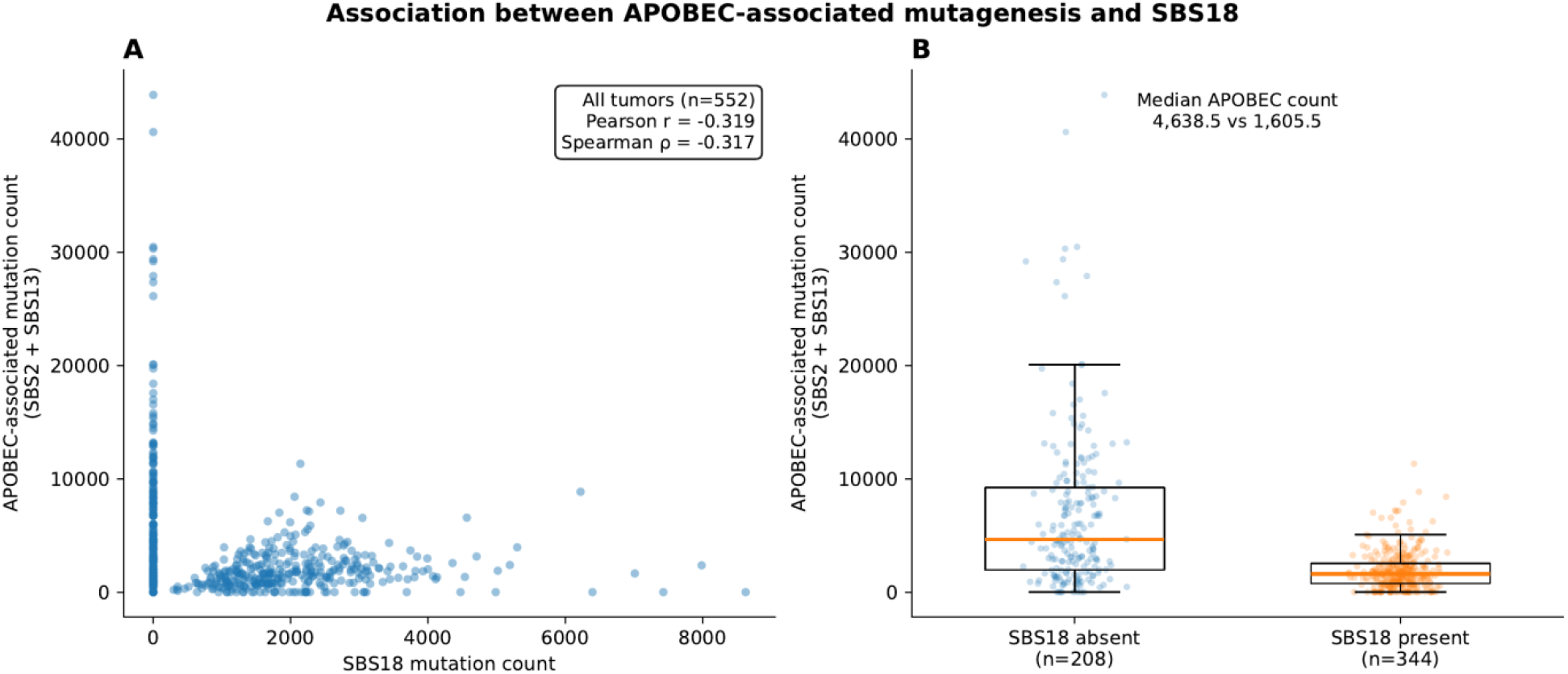
Association between APOBEC-associated mutagenesis and SBS18. (A) Absolute APOBEC mutation count versus SBS18 mutation count across all 552 tumors (absolute count versus absolute count: Pearson r = −0.319; Spearman ρ = −0.317; partial Pearson r = −0.290 after adjustment for non-APOBEC background burden). (B) APOBEC mutation counts in SBS18-absent (n = 208) versus SBS18-present (n = 344) tumors (median 4,638.5 vs 1,605.5). This cohort-level association is primarily driven by SBS18 presence/absence rather than a continuous dose-response relationship among SBS18-positive tumors (see Results).

### Multivariable genomic associations with APOBEC proportion

In the multivariable model, total SBS burden remained positively associated with APOBEC proportion (β = 1.56 × 10□□, P = 0.024), whereas SBS18 (β = −5.19 × 10□□, P = 3.27 × 10□^1^□) and SBS3 (β = −2.17 × 10□□, P = 0.025) showed inverse associations. SBS1 (P = 0.660) and SBS5 (P = 0.135) were not independently associated with APOBEC proportion (Figure 2). The model explained 40.0% of the observed variation (adjusted R^2^ = 0.395). Model diagnostics provided little evidence of problematic multicollinearity, with variance inflation factors ranging from 1.09 to 2.34 and a maximum condition index of 2.89. In 10-fold cross-validation, the mean R^2^ was 0.273 and the median was 0.507, with substantial between-fold variability (SD = 0.576). Bootstrap resampling produced a mean R^2^ of 0.424, with a 95% interval of 0.277–0.595 (Supplementary Table S3).

### Geographic variation in APOBEC-associated mutagenesis

APOBEC proportion differed across the eight countries in unadjusted analysis (Kruskal–Wallis H = 18.59, P = 0.0096; Figure 5A). Country sample sizes were unequal, ranging from seven tumors from the United Kingdom to 178 from Iran. However, after adjustment for total SBS burden and selected mutational signatures, addition of country explained little additional variation in APOBEC proportion (ΔR^2^ = 0.008; F(7,539) = 1.04, P = 0.402), and none of the individual country coefficients had a 95% confidence interval excluding zero (Figure 5B). PCA visualization likewise showed substantial overlap among tumors from different countries (Supplementary Figure S2). These results indicate that the unadjusted geographic heterogeneity was not accompanied by a strong independent country-level association in the adjusted model.

**Fig. 5.**
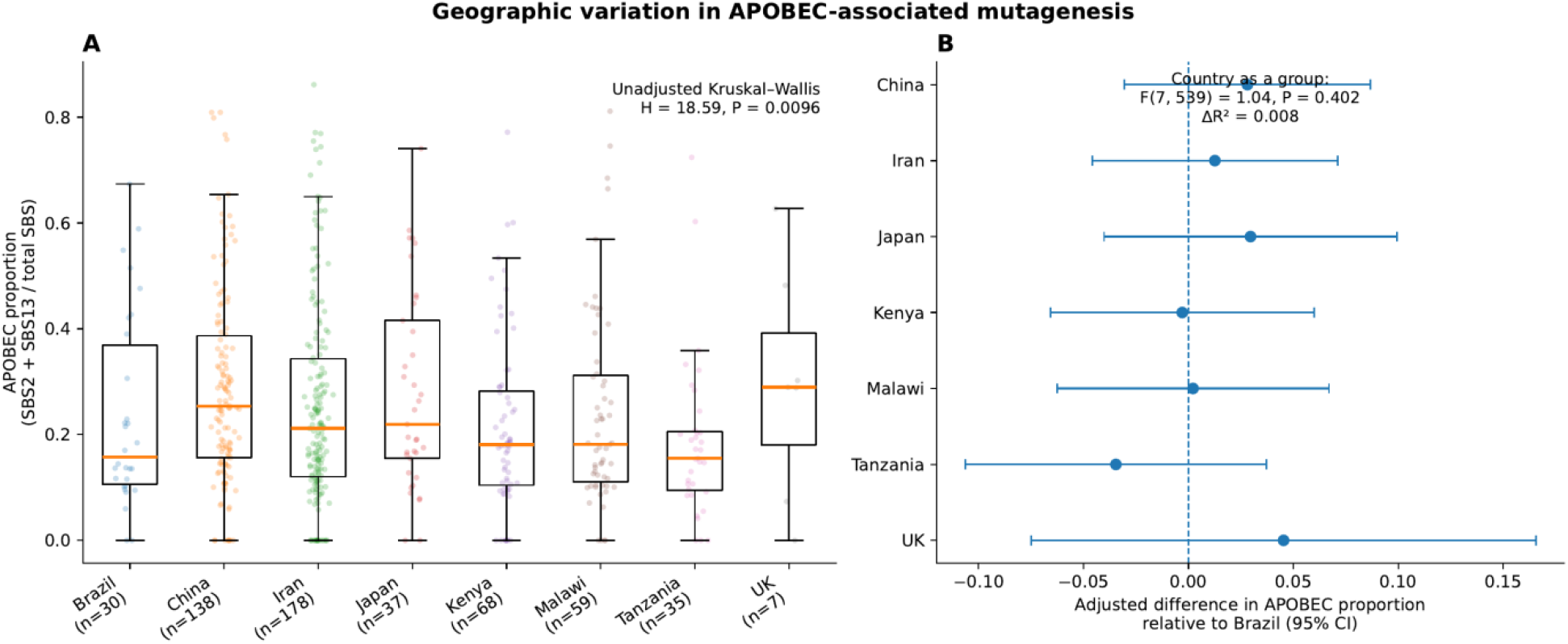
Geographic variation in APOBEC-associated mutagenesis. (A) APOBEC proportion by country (unadjusted Kruskal–Wallis H = 18.59, P = 0.0096). (B) Country coefficients (relative to Brazil) from the genomic-feature-adjusted regression model; country was not significant as a group (F[7,539] = 1.04, P = 0.402; ΔR^2^ = 0.008).

### Driver-gene associations with APOBEC-associated mutagenesis

Analysis of 11 recurrently altered driver genes identified the strongest and most consistent association for *ZNF750* (Figure 6A; Table 3; Supplementary Table S4). *ZNF750*-mutant tumors (n = 12) had a median absolute APOBEC burden of 7,385.5 mutations compared with 1,976 mutations among *ZNF750*-wild-type tumors (n = 540), corresponding to a rank-biserial correlation of 0.610 (Mann–Whitney P = 2.95 × 10□□; FDR-adjusted q = 0.0032; Figure 6B; Table 3). The *ZNF750* association remained significant in the background-adjusted analysis (β = 1.751, 95% CI 0.937–2.565; FDR-adjusted q = 2.73 × 10□□; Table 3; Supplementary Table S4). *KMT2D* (q = 0.0091) and *PIK3CA* (q = 0.0226) were significant only in the rank-based analysis, whereas *AJUBA* was significant only in the adjusted analysis (q = 0.0280). Thus, *ZNF750* was the only gene supported by both analytical approaches. Among the 12 *ZNF750*-mutant tumors, nine carried nonsense variants and three carried missense variants. Only two of the 12 variants occurred in canonical SBS2/SBS13 sequence contexts. Given the small number of *ZNF750*-mutant tumors and the observational nature of these analyses, this association should be considered hypothesis-generating rather than evidence of a direct mechanistic relationship.

**Table 3.** Associations between driver-gene alterations and absolute APOBEC-associated mutation burden across all 11 prespecified ESCC driver genes, with Benjamini–Hochberg FDR correction (rank-based and background-burden-adjusted analyses).

| **Gene** | **Mutant n (%)** | **Median APOBEC (mut. vs WT)** | **Rank-biserial r** | **MW P** | **MW BH-FDR q** | **Adj. β (95% CI)** | **Adj. P** | **Adj. BH-FDR q** | **Evidence pattern** |
|---|---|---|---|---|---|---|---|---|---|
| ZNF750 | 12 (2.2%) | 7385.500 vs 1976.000 | 0.610 | 2.95e-04 | 0.003 | 1.751 (0.937 to 2.565) | 2.48e-05 | 2.73e-04 | Supported in both analyses |
| KMT2D | 69 (12.5%) | 3678.000 vs 1942.000 | 0.234 | 0.002 | 0.009 | 0.296 (-0.374 to 0.967) | 0.386 | 0.744 | Rank-based analysis only |
| PIK3CA | 48 (8.7%) | 3733.000 vs 1960.500 | 0.239 | 0.006 | 0.023 | 0.366 (-0.429 to 1.161) | 0.366 | 0.744 | Rank-based analysis only |
| FAT1 | 39 (7.1%) | 2723.000 vs 1976.000 | 0.175 | 0.069 | 0.189 | 0.074 (-0.878 to 1.027) | 0.879 | 0.959 | Not FDR-significant |
| NOTCH1 | 53 (9.6%) | 1842.000 vs 2026.000 | -0.122 | 0.144 | 0.281 | -0.577 (-1.361 to 0.208) | 0.150 | 0.550 | Not FDR-significant |
| KEAP1 | 14 (2.5%) | 3733.000 vs 1995.000 | 0.223 | 0.153 | 0.281 | 0.433 (-0.930 to 1.795) | 0.534 | 0.744 | Not FDR-significant |
| AJUBA | 10 (1.8%) | 2562.500 vs 1997.500 | 0.180 | 0.330 | 0.518 | 0.818 (0.245 to 1.390) | 0.005 | 0.028 | Adjusted log-scale analysis only |
| NFE2L2 | 35 (6.3%) | 2253.000 vs 1995.000 | 0.037 | 0.714 | 0.872 | 0.224 (-0.495 to 0.943) | 0.541 | 0.744 | Not FDR-significant |
| TP53 | 420 (76.1%) | 2056.000 vs 1867.000 | 0.025 | 0.665 | 0.872 | -0.032 (-0.494 to 0.429) | 0.891 | 0.959 | Not FDR-significant |
| CDKN2A | 62 (11.2%) | 2454.000 vs 1997.500 | 0.013 | 0.866 | 0.953 | -0.224 (-0.913 to 0.465) | 0.524 | 0.744 | Not FDR-significant |
| CUL3 | 11 (2.0%) | 1861.000 vs 2012.000 | -0.007 | 0.970 | 0.970 | -0.043 (-1.668 to 1.582) | 0.959 | 0.959 | Not FDR-significant |

**Fig. 6.**
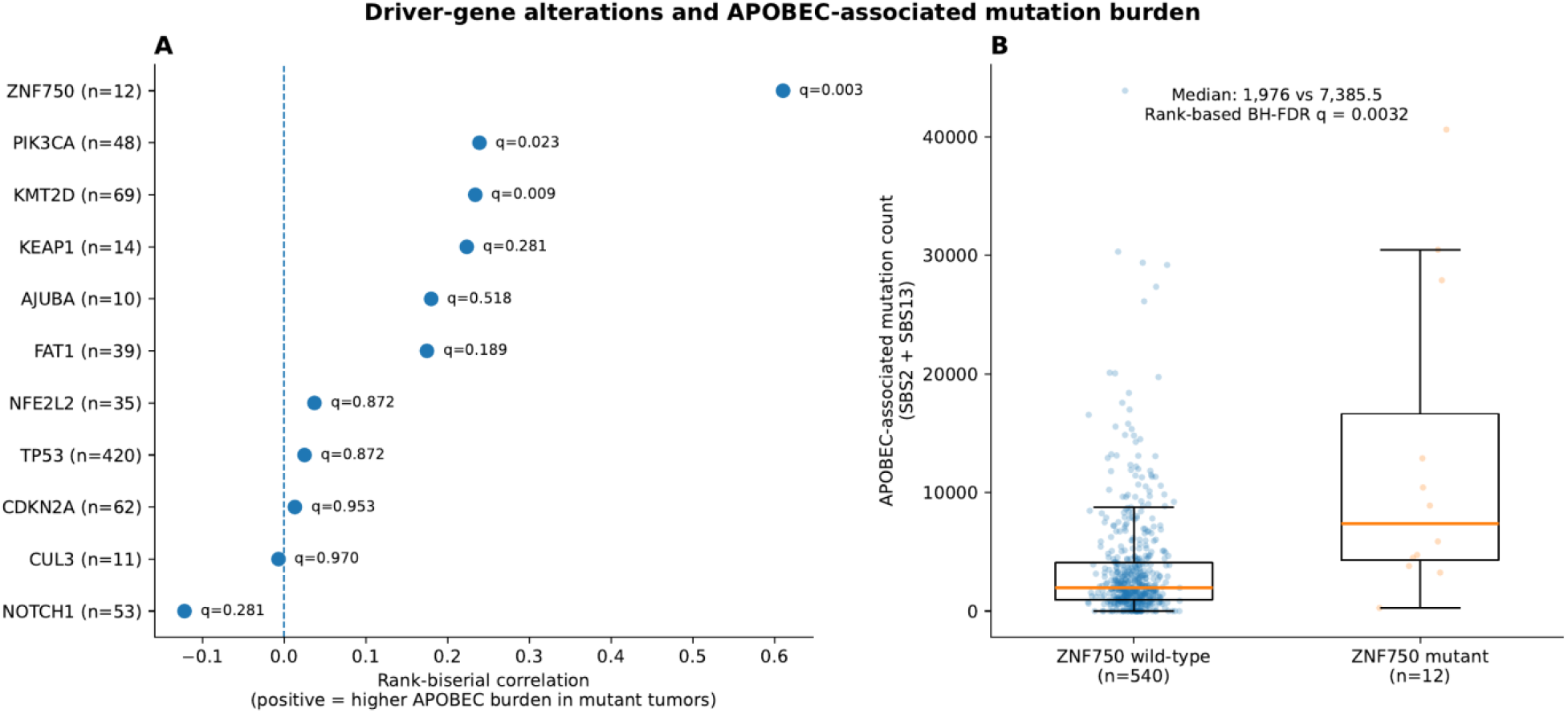
Driver-gene alterations and APOBEC-associated mutation burden. (A) Rank-biserial correlation and BH-FDR q values for all 11 prespecified driver genes (positive = higher APOBEC burden in mutant tumors). (B) APOBEC mutation counts in ZNF750 wild-type (n = 540) versus ZNF750-mutant (n = 12) tumors (median 1,976 vs 7,385.5; BH-FDR q = 0.0032).

## Discussion

In this study, we examined genomic correlates of intertumoral variation in APOBEC-associated mutagenesis across 552 ESCC tumors from eight countries. Three findings emerged from the revised analyses. First, several relationships observed when mutational signatures were expressed as proportions were sensitive to the compositional nature of the data and did not persist when absolute mutation counts were considered. Second, SBS18 was an important exception: its inverse association with APOBEC-associated mutagenesis remained evident in absolute-count analyses, although further analysis showed that this cohort-level relationship was largely characterized by differences between tumors with and without detectable SBS18 rather than by a continuous inverse relationship among SBS18-positive tumors. Third, ZNF750 alteration was associated with higher absolute APOBEC burden across two complementary analytical approaches, although the small number of ZNF750-mutant tumors requires cautious interpretation. Together, these findings refine the genomic context of APOBEC-associated mutagenesis in ESCC while emphasizing the importance of accounting for the statistical structure of mutational signature data.

APOBEC-associated mutagenesis is already recognized as a prominent component of the ESCC mutational landscape [6,7]. In the international cohort on which the present secondary analysis is based, Moody et al. reported widespread SBS2 and SBS13 activity across tumors from countries with markedly different ESCC incidence [5]. Our analysis therefore does not establish the presence or geographic distribution of APOBEC mutagenesis in ESCC. Rather, it examines why the relative and absolute contributions of APOBEC-associated mutations vary among tumors within this cohort and evaluates whether associations with other genomic features remain robust to alternative representations of mutational signature activity.

This distinction is also important when considering the term “determinants” of APOBEC mutagenesis. Previous studies have identified molecular factors more directly related to APOBEC enzymatic activity. For example, a germline copy-number polymorphism involving *APOBEC3A* and *APOBEC3B* has been associated with increased APOBEC-associated mutation burden in cancer [12], while experimental studies have demonstrated direct mutagenic contributions of APOBEC3A and APOBEC3B and differences in their sequence preferences and relative activities [14]. Other work has provided evidence that APOBEC3A is a major contributor to APOBEC-associated mutation in several cancer contexts [8]. These germline, transcriptional, and enzyme-specific factors were not directly measured in the present analysis. Accordingly, the genomic features identified here are more appropriately considered correlates of APOBEC-associated mutagenesis rather than molecular determinants.

An important methodological finding was that several apparent relationships between APOBEC-associated mutagenesis and other signatures depended strongly on whether signature activity was represented as a proportion or an absolute mutation count. Mutational signature proportions are compositional [15–17]: increasing the proportional contribution of one signature necessarily reduces the relative contribution available to others even when their absolute mutation counts do not decrease. Consistent with this constraint, SBS1, SBS3, and SBS5 showed inverse relationships with APOBEC-associated mutagenesis in proportional analyses but positive relationships when absolute mutation counts were examined. These direction reversals indicate that the proportional associations should not be interpreted as evidence that increasing APOBEC proportion suppresses these mutational processes. They also illustrate why biological interpretations based exclusively on relative signature contributions can be misleading when one process accounts for a substantial fraction of the total mutation burden.

SBS18 differed from this pattern. The inverse relationship between APOBEC-associated mutagenesis and SBS18 weakened in magnitude when absolute rather than proportional activities were examined but remained evident using both Pearson and rank-based correlations (proportion versus proportion, ρ = −0.527; absolute count versus absolute count, ρ = −0.317; partial Pearson r = −0.290 after adjustment for non-APOBEC background burden). This suggests that the association cannot be explained entirely by the constant-sum constraint of proportional signature data. Nevertheless, further analysis substantially changed its biological interpretation. Tumors without detectable SBS18 had considerably higher APOBEC mutation burdens than tumors in which SBS18 was detected. However, among SBS18-positive tumors, there was no evidence of a continuous inverse relationship after accounting for background mutation burden. The overall inverse association is therefore better characterized as a cohort-level difference associated with SBS18 presence or absence than as evidence for progressive antagonism between APOBEC-mediated and SBS18-associated mutagenesis.

SBS18 has been associated with oxidative DNA damage, particularly damage related to reactive oxygen species, and this signature is notably common in inflammation-associated upper gastrointestinal tumors including esophageal and gastric adenocarcinoma [18]. APOBEC-associated mutagenesis, in contrast, results from cytidine deamination in single-stranded DNA substrates and can be promoted by cellular states that increase exposure of single-stranded DNA during replication, transcription, or DNA repair [19]. These mechanistic differences provide biological context for the observed heterogeneity, but our data do not establish that the two processes compete with or suppress one another. The previous interpretation of a distinct APOBEC-high state characterized by reduced oxidative mutagenesis is therefore too strong. Instead, the present findings identify an association between APOBEC burden and SBS18 status that warrants investigation in datasets containing direct measures of oxidative stress, DNA repair activity, APOBEC expression, and tumor evolutionary history.

The PCA sensitivity analysis provides a related example of the importance of separating descriptive structure from independent evidence. In the original PCA, PC1 was strongly associated with APOBEC proportion, but SBS13 and SBS2 were themselves the largest contributors to PC1. When SBS2 and SBS13 were removed before PCA, the association between APOBEC proportion and PC1 disappeared. The original PCA therefore primarily describes the contribution of APOBEC-associated signatures to overall mutational variation rather than identifying an independent genomic axis associated with APOBEC-associated mutagenesis. We consequently regard PCA as a descriptive component of the analysis rather than evidence for a separate APOBEC-associated tumor state.

The multivariable model similarly requires cautious interpretation. Although the model accounted for approximately 40% of APOBEC-proportion variation within the analyzed cohort, internal cross-validation showed lower and highly variable performance across held-out subsets. Conventional diagnostics did not indicate substantial numerical multicollinearity, but coefficient-stability analyses showed that estimates for several signatures were sensitive to model specification. Moreover, modeling a proportional APOBEC outcome together with total mutation burden and component signature activities retains an inherent part-whole relationship that variance inflation factors cannot resolve. The model therefore summarizes associations within this dataset and should not be interpreted as a validated predictive model or as evidence that the included variables mechanistically determine APOBEC proportion.

The driver-gene analysis identified *ZNF750* alteration as a potentially interesting correlate of APOBEC-associated mutation burden. *ZNF750*-mutant tumors had higher absolute APOBEC counts than wild-type tumors, and this association remained after adjustment for non-APOBEC background mutation burden and correction for testing 11 driver genes. Importantly, only a minority of the observed *ZNF750* variants occurred in sequence contexts compatible with canonical SBS2 or SBS13 mutagenesis, arguing against the simple explanation that the tumor-level association arose because most *ZNF750* alterations were themselves direct APOBEC-context mutations, although recurrent mutations in APOBEC-high tumors can also arise as passenger hotspots independent of driver selection [20]. Nevertheless, only 12 tumors carried protein-altering *ZNF750* mutations. The estimate is therefore based on a small subgroup and requires validation in an independent ESCC cohort. The less consistent results for *KMT2D, PIK3CA*, and *AJUBA*, which depended on the analytical approach used, further support treating the driver-gene analysis as hypothesis-generating rather than evidence of causal relationships.

Geographic analyses also require a more qualified interpretation than suggested by the original analysis. APOBEC proportions differed across countries in the unadjusted comparison, but addition of country to the genomic regression model produced only a small increase in model R^2^, and country was not statistically significant when the seven indicator variables were tested jointly. This indicates that country contributed little additional information after accounting for the genomic variables included in that particular model. It does not establish that geographic, environmental, or population-associated factors are unrelated to APOBEC mutagenesis. Indeed, the source ESCC study identified associations between particular mutational signatures and exposures or germline variation, demonstrating that epidemiological and inherited factors can contribute to mutational patterns in this cohort [5]. Country is also an imperfect proxy for these individual-level exposures.

Several limitations are important when interpreting these findings. First, this was a secondary observational analysis of previously generated whole-genome sequencing data, precluding causal inference or direct experimental evaluation of the mechanisms underlying the observed associations. Second, mutational signature proportions are compositional, creating dependencies among signatures. Although absolute-count sensitivity analyses substantially improved interpretation, absolute mutation counts have their own dependence on overall mutation burden and sequencing-derived mutation detection. Third, individual-level covariates that could confound geographic and genomic associations—including age, tobacco use, alcohol consumption, and tumor purity—were not available in the metadata used for the present analysis, and biological sex was unavailable for most tumors. The country groups were also markedly unbalanced, with particularly small representation from some populations. Fourth, the multivariable model was internally evaluated but not externally validated, and its cross-validated performance was variable. Fifth, several driver-gene subgroups were small, particularly the *ZNF750*-mutant group, increasing uncertainty around gene-specific estimates. Finally, this analysis did not include direct measurements of *APOBEC3A* or *APOBEC3B* expression or activity, germline APOBEC variation, epigenetic regulation, tumor microenvironment, or temporal and intratumoral heterogeneity. These factors may account for substantial additional variation in APOBEC-associated mutagenesis.

Despite these limitations, the study has several strengths. The analysis uses a large whole-genome ESCC cohort spanning eight countries and evaluates APOBEC-associated mutagenesis as a continuous phenotype. More importantly, the revised analytical framework directly tests the robustness of associations across proportional and absolute representations, evaluates PCA circularity, examines model stability and internal performance, applies multiple-testing correction across prespecified driver genes, and distinguishes exploratory associations from more consistent findings. This approach provides a more conservative characterization of genomic correlates of APOBEC-associated mutagenesis than would be obtained from proportional signature analyses alone.

Future work should evaluate the APOBEC–SBS18 and APOBEC—*ZNF750* relationships in independent ESCC cohorts and integrate genomic signatures with direct molecular measurements. In particular, combining whole-genome sequencing with *APOBEC3A* and *APOBEC3B* expression and activity, germline APOBEC variation, oxidative-stress markers, DNA repair phenotypes, and longitudinal or multiregion sequencing could help distinguish factors that precede APOBEC activation from genomic changes that accumulate as a consequence of it. Such studies will be necessary to move from statistical correlates toward mechanistic determinants of APOBEC-associated mutagenesis in ESCC.

## Conclusions

APOBEC-associated mutagenesis varies substantially among ESCC tumors, but the apparent relationships between APOBEC-associated mutagenesis and other mutational signatures depend in part on how mutational composition is represented. Inverse associations involving SBS1, SBS3, and SBS5 were not maintained in absolute-count analyses, whereas the inverse relationship with SBS18 persisted at the cohort level. Further analysis indicated that this APOBEC–SBS18 relationship was primarily characterized by differences between tumors with and without detectable SBS18 rather than by a continuous inverse relationship among SBS18-positive tumors. *ZNF750* alteration was also associated with increased APOBEC mutation burden across complementary analyses, although the small number of mutant tumors makes this finding hypothesis-generating. Country contributed little additional variation after accounting for the genomic variables included in the model, but the available data do not exclude geographic or exposure-related influences. Overall, these findings identify genomic correlates of APOBEC-associated mutagenesis in ESCC while demonstrating the importance of compositional sensitivity analyses and cautious distinction between statistical associations and biological determinants.

## Acknowledgements

The authors acknowledge the investigators of the original international ESCC whole-genome sequencing studies for generating and making the datasets available through the European Genome–Phenome Archive.

## Authors’ contributions

EOA conceived and designed the study, developed the methodology, performed the formal analyses, generated the visualizations, interpreted the findings, and wrote the original draft of the manuscript. JAY contributed to methodology, data interpretation, and critical revision of the manuscript. EYK contributed to study supervision, scientific interpretation, and critical review of the manuscript. POM, KF, IKA, and EAB contributed to the review and editing of the manuscript. All authors have read and approved the final version of the manuscript. Emmanuel Owusu Ansah, the corresponding author and manuscript guarantor, had full access to all study data and takes responsibility for the integrity of the data and the accuracy of the analyses.

## Transparency declaration

The lead author and manuscript guarantor, Emmanuel Owusu Ansah, affirms that this manuscript is an honest, accurate, and transparent account of the study being reported; that no important aspects of the study have been omitted; and that any discrepancies from the study as planned have been explained.

## ORCID

Emmanuel Owusu Ansah: https://orcid.org/0000-0003-4958-0934

## Availability of data and materials

The whole-genome sequencing data analyzed in this study were obtained from the European Genome–Phenome Archive (EGA) under accession number EGAD00001006868 (BAM files), associated with the published international ESCC cohort study [5]. Filtered somatic single-nucleotide variant calls used for mutational signature analysis were obtained from the companion dataset EGAD00001008339, deposited under the same parent study (EGAS00001002725). Owing to controlled-access restrictions, raw sequencing data are available from the EGA upon reasonable request and approval by the relevant data access committee. Processed data generated during the current study, including mutational signature summaries and statistical outputs, are provided within the article and its supplementary information files.

## Code availability

The Python scripts used for the statistical, sensitivity, and supplementary analyses reported in this study are publicly available at https://github.com/Nana1995365/ESCC_APOBEC_Reproducibility. The repository includes dependency specifications and instructions for executing the analyses. The underlying sample-level genomic datasets are not included in the repository and remain subject to the applicable data-access and sharing conditions.

